# Finding A Fresh Carcass: Bacterially-Derived Volatiles And Burying Beetle Search Success

**DOI:** 10.1101/2020.01.25.919696

**Authors:** Stephen T. Trumbo, Sandra Steiger

**Affiliations:** University of Connecticut, Department of Ecology and Evolutionary Biology, Waterbury, Connecticut, USA; University of Bayreuth, Department of Evolutionary Animal Ecology, Bayreuth, Germany

**Keywords:** Carrion ecology, Methyl thiocyanate, Dimethyl trisulfide, Forensic entomology, *Nicrophorus*, Semiochemical

## Abstract

When burying beetles first emerge as adults, they search for well-rotted carcasses with fly maggots on which to feed. After attaining reproductive competence, they switch their search and respond to a small, fresh carcass to prepare for their brood. Because the cues used to locate a feeding versus a breeding resource both originate from carrion, the beetles must respond to subtle changes in volatiles during decomposition. We investigated cues used to locate a fresh carcass in the field by (1) a general subtractive method, applying an antibacterial or antifungal to reduce volatiles, and (2) a specific additive method, placing chemicals near a fresh carcass. Five sulfur-containing compounds were studied: dimethyl sulfide (DMS), dimethyl disulfide (DMDS), dimethyl trisulfide (DMTS), methyl thiolacetate (MeSAc) and methyl thiocyanate (MeSCN). For the sulfides, we predicted that DMS would be the most attractive and DMTS the least attractive because of differences in the timing of peak production. We made no *a priori* predictions for MeSAc and MeSCN. Antibacterial treatment of a carcass aged for 48 h resulted in a 59% decrease in beetles discovering the resource. The addition of MsSAc had no effect on discovery of a fresh carcass, while DMS and DMDS had a limited ability to attract breeding beetles. The chemical that was least well known, MeSCN, had a remarkable effect, increasing beetle numbers by 200-800% on a fresh carcass and almost guaranteeing discovery. DMTS, which is known to attract a variety of carrion insects, was the only compound to significantly reduce beetle presence at a fresh carcass. A laboratory experiment demonstrated that DMTS does not directly inhibit breeding, suggesting that DMTS deters breeding beetles while they fly.

## INTRODUCTION

A complex life requires responses to a series of different cues to organize activities such as feeding, shelter-seeking and reproduction. This has been appreciated since von Uexküll’s classic work described the sequence of signs that allow a tick to locate and exploit its host (see Agamben 2004). The changes in cue-response can be both dramatic and rapid. The seabird tick (*Ixodes uriae* White) attends to very different volatiles to move from conspecific aggregations to their host (Benoit et al. 2008), while the green bottle fly, *Lucilia sericata* Meigen, responds to fecal cues for feeding but to carrion cues for breeding (Brodie et al. 2016). The burying beetles (*Nicrophorus* spp.) likewise use different resources for feeding versus reproduction (von Hoermann et al. 2013). In their case, however, the distinctive cues must be subtle because they all originate from a dead animal, albeit from different successional stages. The beetles must either attend to compounds that are prominent at different successional stages or to changing proportions of compounds. Investigating the burying beetle umwelt is further complicated because over 500 volatiles have been tabulated for animal decomposition (Cammack et al. 2015; Forbes and Carter 2015). Identifying the changing cues used by insects during decomposition will be helpful to understand how critical nutrients move through the ecosystem and the succession of insects on a corpse, the later of which has forensic applications (Merritt and De Jong 2015). There is a particular gap in our knowledge of the cues used by early colonizers of a carcass, when the volatile profile is barely distinguishable from a living organism (Armstrong et al. 2016; Tomberlin et al. 2011).

Part of the story of burying beetle responses to cues is known. When female beetles first emerge as adults, they, like mosquitoes, do not breed until sufficient feeding allows the ovaries to increase in size and then plateau, waiting for a reproductive cue (Trumbo et al. 1995; Trumbo 1997). During the one to three week feeding period they will come to a carcass of any size, preferring ones in active decay where they consume carrion and fly maggots. Von Hoermann et al. (2013) speculate that newly emerged beetles (or beetles in reproductive diapause) avoid a fresh carcass and thereby avoid breeding congeners that will fight, sometimes to the death — a high fitness cost for a non-breeder that just wants a meal. At reproductive maturity, a burying beetle searches for a new type of resource, a small, not-too-decomposed vertebrate carcass that will stimulate final ovarian maturation (Wilson and Knollenberg 1984). They will bury the appropriate carcass and prepare it for their brood underground (Pukowski 1933). Although their reproductive success is highest on the freshest carcasses (Rozen et al. 2008; Trumbo et al. 2016), they have some difficulty locating an animal that has been dead for less than 24 h, presumably because of the scarcity of distinguishing cues (Trumbo 2016). Burying beetles (nicrophorine silphids), which evolved at least 125 mya (Cai et al. 2014; Sikes and Venables 2013), have been making the distinction between feeding and breeding resources for a long time, as their ancestors, like modern, less parental silphine carrion beetles, likely used older carcasses for both feeding and reproduction (Anderson and Peck 1985; Ratcliffe 1996).

Burying beetles, in common with other carrion insects, are highly sensitive to sulfur-containing volatile organic compounds (S-VOCs; Fig. 1) (Kalinova et al. 2009) that are produced by microbes on carrion (Cernosek et al. 2020; Crippen et al. 2015; Stutz et al. 1991). The sulfides, thought to be the most important cues for carrion insects (Kalinova et al. 2009, reviewed in Cammack et al. 2015), form a series from the most volatile (DMS) which has the earliest production peak on a decomposing mouse to the least volatile, dimethyl tetrasulfide (DMQS), which has a later peak.

**Fig. 1.**
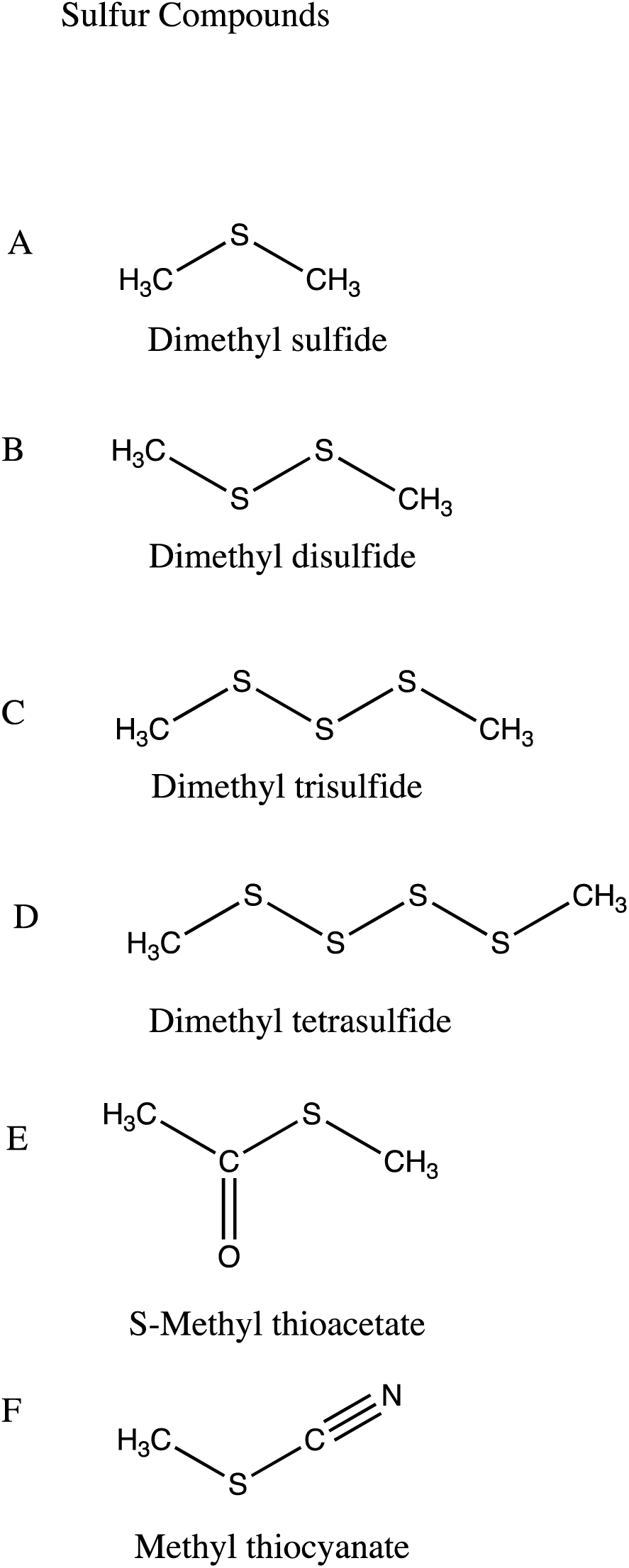
Six sulfur-containing volatile organic compounds released from carrion (S-methyl thioacetate = methyl thiolacetate)

Dimethyl disulfide (DMDS) and dimethyl trisulfide (DMTS) are intermediate in volatility and timing of peak production (Kalinova et al. 2009). DMDS and DMTS have also been identified as critical volatiles from corpse-mimicking plants that exploit deceived pollinators (Jürgens and Shuttleworth 2015). DMS, DMDS and especially DMTS are clearly attractive to burying beetles (Podskalska et al. 2009) although the life stage of attracted beetles was not clear. Methyl thiolacetate (MeSAc), also produced by corpse-mimicking plants, elicits electrical activity in isolated antennae of burying beetles (Kalinova et al. 2009); it has not been the subject of behavioral assays. We know of no prior work on methyl thiocyanate (MeSCN) in insects.

In the present study, we investigated the importance of microbes and five microbially-derived S-VOCs for inducing free-flying burying beetles to locate and bury a fresh carcass in the field. We first employed a general subtractive method (antibacterial and antifungal treatment of a carcass) and then a specific additive method (chemical supplements near a carcass). The supplements assayed were DMS, DMDS, DMTS, MeSAc and MeSCN. Based on the work on the sulfides by Kalinova et al. (2009), we predicted that all sulfides would be attractive, with DMS, the earliest to peak in production, the most attractive and DMTS the least. We made no *a priori* predictions for MeSAc and MeSCN, for which there is little background.

## METHODS AND MATERIALS

### General Methods for Field Experiments

Seven field experiments, similar in design, were conducted to examine whether bacteria are an important source of cues for burying beetles attempting to locate a carcass for breeding (Exps. 1 and S1) and whether known microbially-derived S-VOCs attract or repel burying beetles to a fresh carcass (Exps. 2-6). Three secondary forests were used to minimize vertebrate scavenging and to allow concurrent experiments without interference (Bethany, USA 41^0^27^1^36^11^N, 72^0^57^1^37^11^W; Woodbury, USA 41^0^31^1^48^11^N, 73^0^10^1^12^11^W; Colebrook, USA 42^0^00^1^08^11^N, 73^0^04^1^36^11^W).

At each site, three-point transects were established with transect points separated by 20 m (the DMTS experiment employed a 6-point transect). The number of transects employed (2 or 3) depended on whether one or two treatments were being tested versus a control. A single transect consisted of multiple carcasses of the same treatment, and transects were greater than 200 m apart to reduce cross-attraction between transect-treatments. In all experiments, free-flying beetles had an opportunity to discover and bury a mouse carcass in a cup to measure breeding activity. Cups (10 cm diameter, 12 cm height) were 4/5^th^filled with soil from the field and buried in the ground so that the rim of the cup was flush with the ground surface. A recently thawed mouse carcass (8 – 12 g, Rodent Pro**®**, Inglefield, IN, U.S.A) was placed on top of the soil in the cup. To test chemical supplements, a microcentrifuge tube (1.5. ml, 4 cm height) with a hole made by a hypodermic needle (26 g for the more volatile DMS, 23 g for all other chemicals, Exelint) was placed on top of the soil in the cup with enough chemical to last the duration of the trial on the warmest expected days. When the most active burying beetle was nocturnal (*N. orbicollis* Say; trials between 1 June and 24 August), carcasses were placed in the field at 17:00 and checked at 9:00 the following day. When the most active burying beetle was diurnal (*N. tomentosus* Weber; 25 August – 25 September), carcasses were placed in the field at 10:00 and checked the same day at 18:00. A trial was scored as a successful discovery by a breeder if the carcass was buried in the cup and beetles were present. A carcass that was removed from the site was scored as vertebrate scavenging. A carcass that remained on top of the soil was scored as not discovered. After each trial, all cups were returned to the laboratory for cleaning to remove residual odor and non-volatized chemical was stored (-7°C) for later use. Treatments were rotated through the transects such that each transect was used once for each treatment before a transect was re-used for the same treatment. In this way, each transect was used an equal number of times for each treatment, minimizing location and location x season biases.

### Experiment 1 – Antimicrobial Treatment of a Carcass

To examine whether bacteria are the source of cues that beetles use to discover resources, carcasses of three types (fresh, aged 2 days or aged 2 days with antibacterial treatment) were placed in the field (Bethany site). Fresh carcasses were thawed on soil at room temperature for 2-4 h and then immersed in water and gently tumbled for 1 min in a closed container, prior to placement in the field. Two-day carcasses were thawed and aged in the laboratory for 48 h with water immersions at 3 h, 24 h and 48 h. Between immersions, carcasses were placed on soil from the woodland to dry and to expose them to naturally-occurring soil bacteria. Antibacterial-treated carcasses were immersed in an aqueous tetracycline hydrochloride solution (45 mg per 100ml) (Sigma Aldrich) and tumbled to ensure complete topical exposure at 3 h, 24 h and 48 h, with placement on soil in between. Nine mouse carcasses (3 for each of the 3 treatments) were placed in the field on each of 18 dates (3 August – 24 September 2010, 19 June – 25 September 2013) (total of 162 carcasses, 54 per treatment).

### Experiments 2-6 – Chemical Supplements

Five sulfur-containing compounds, the three sulfides, DMS, DMDS and DMTS, and MeSAc and MeSCN (Sigma Aldrich) were tested for their effects on attracting breeding burying beetles to a fresh carcass, in a similar manner as described above. In each test of sulfur-based compounds, a fresh carcass was used as the control treatment and a fresh carcass with an adjacent microcentrifuge tube containing chemical was the experimental treatment. The locations of trials, amount of chemical, sample sizes and target species for Experiments 2–6 are shown in Table 1. DMS was tested on 12 dates in midsummer (15 August – 30 August 2016, 5 June – 12 June 2019) and 12 dates in late summer (12 September – 19 September 2016, 13 September – 20 September 2019) (Exps. 2a & 2b). DMDS was tested on 16 dates in midsummer (14 July – 10 August 2016, 2 June – 12 July 2017) (Exp. 3). DMTS was tested on six dates in midsummer (10 July – 25 July 2019) (Exp. 4).

**Table 1.**
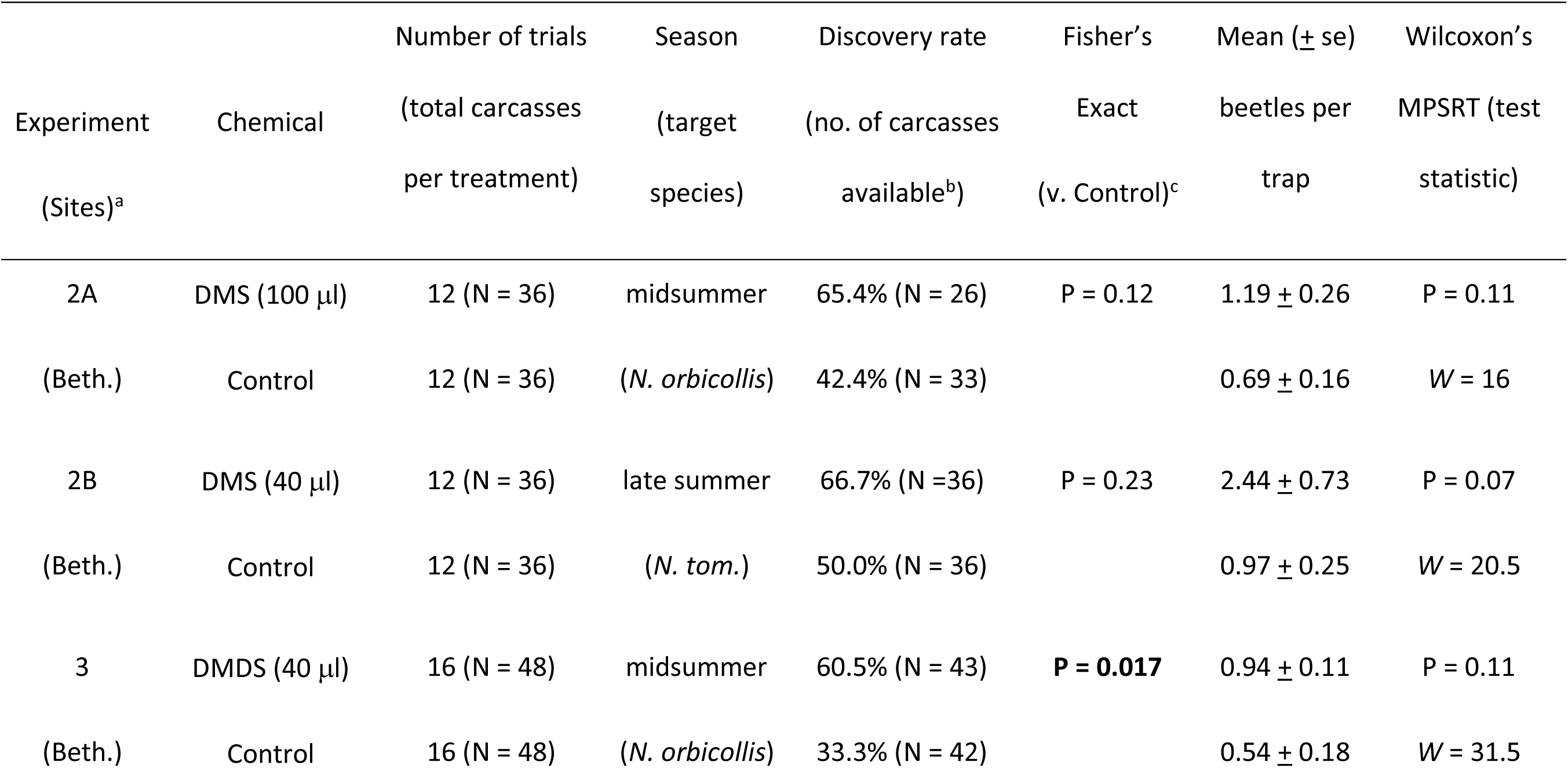
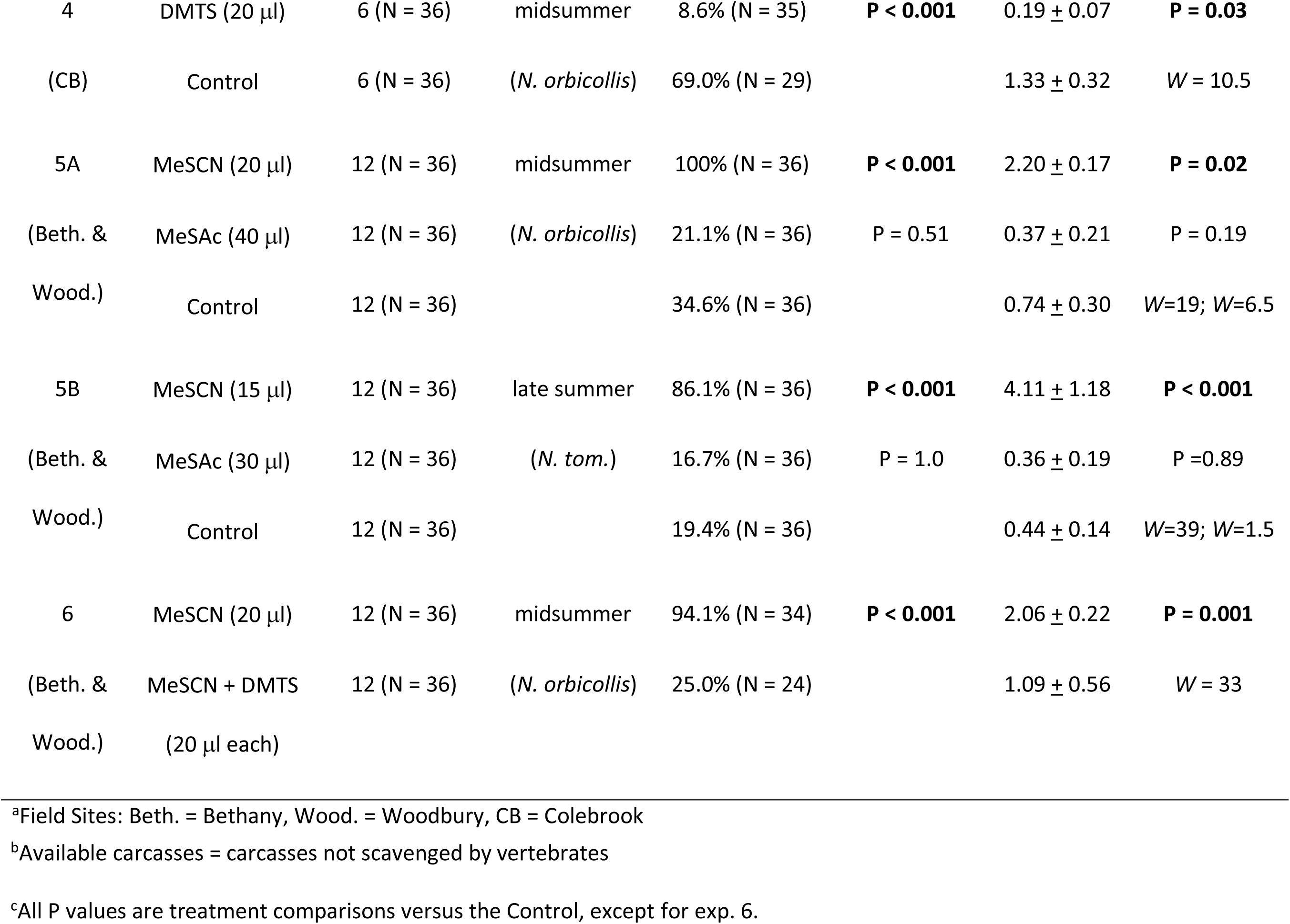
Experiments to evaluate the effects of chemical supplements on attraction of *N. orbicollis* and *N. tomentosus* to a fresh carcass

| Experiment<br>(Sites)<sup>a</sup> | Chemical | Number of trials<br>(total carcasses<br>per treatment) | Season<br>(target<br>species) | Discovery rate<br>(no. of carcasses<br>available<sup>b</sup>) | Fisher's<br>Exact<br>(v. Control)<sup>c</sup> | Mean (± se)<br>beetles per<br>trap | Wilcoxon's<br>MPSRT (test<br>statistic) |
| --- | --- | --- | --- | --- | --- | --- | --- |
| 2A<br>(Beth.) | DMS (100 μl)<br>Control | 12 (N = 36)<br>12 (N = 36) | midsummer<br>(*N. orbicollis*) | 65.4% (N = 26)<br>42.4% (N = 33) | P = 0.12 | 1.19 ± 0.26<br>0.69 ± 0.16 | P = 0.11<br>W = 16 |
| 2B<br>(Beth.) | DMS (40 μl)<br>Control | 12 (N = 36)<br>12 (N = 36) | late summer<br>(*N. tom.*) | 66.7% (N = 36)<br>50.0% (N = 36) | P = 0.23 | 2.44 ± 0.73<br>0.97 ± 0.25 | P = 0.07<br>W = 20.5 |
| 3<br>(Beth.) | DMDS (40 μl)<br>Control | 16 (N = 48)<br>16 (N = 48) | midsummer<br>(*N. orbicollis*) | 60.5% (N = 43)<br>33.3% (N = 42) | **P = 0.017** | 0.94 ± 0.11<br>0.54 ± 0.18 | P = 0.11<br>W = 31.5 |
| 4 | DMTS (20 µl) | 6 (N = 36) | midsummer | 8.6% (N = 35) | **P < 0.001** | 0.19 ± 0.07 | **P = 0.03** |
| (CB) | Control | 6 (N = 36) | (*N. orbicollis*) | 69.0% (N = 29) |  | 1.33 ± 0.32 | W = 10.5 |
| 5A | MeSCN (20 µl) | 12 (N = 36) | midsummer | 100% (N = 36) | **P < 0.001** | 2.20 ± 0.17 | **P = 0.02** |
| (Beth. & | MeSAc (40 µl) | 12 (N = 36) | (*N. orbicollis*) | 21.1% (N = 36) | P = 0.51 | 0.37 ± 0.21 | P = 0.19 |
| Wood.) | Control | 12 (N = 36) |  | 34.6% (N = 36) |  | 0.74 ± 0.30 | W=19; W=6.5 |
| 5B | MeSCN (15 µl) | 12 (N = 36) | late summer | 86.1% (N = 36) | **P < 0.001** | 4.11 ± 1.18 | **P < 0.001** |
| (Beth. & | MeSAc (30 µl) | 12 (N = 36) | (*N. tom.*) | 16.7% (N = 36) | P = 1.0 | 0.36 ± 0.19 | P = 0.89 |
| Wood.) | Control | 12 (N = 36) |  | 19.4% (N = 36) |  | 0.44 ± 0.14 | W=39; W=1.5 |
| 6 | MeSCN (20 µl) | 12 (N = 36) | midsummer | 94.1% (N = 34) | **P < 0.001** | 2.06 ± 0.22 | **P = 0.001** |
| (Beth. & | MeSCN + DMTS | 12 (N = 36) | (*N. orbicollis*) | 25.0% (N = 24) |  | 1.09 ± 0.56 | W = 33 |
| Wood.) | (20 µl each) |  |  |  |  |  |  |

<sup>a</sup>Field Sites: Beth. = Bethany, Wood. = Woodbury, CB = Colebrook
<sup>b</sup>Available carcasses = carcasses not scavenged by vertebrates
<sup>c</sup>All P values are treatment comparisons versus the Control, except for exp. 6.

MeSAc and MeSCN were both tested versus a control in the same experiment on 12 dates in midsummer (22 June – 20 July 2018, Exp. 5a) and 12 dates in late summer (24 August – 19 September 2018, Exp. 5b). Exp. 6 examined whether the identified repellent (DMTS) had a negative effect on discovery when paired with the identified most highly attractive supplement (MeSCN). A fresh carcass was supplemented with either MeSCN or a combination of MESCN and DMTS (in separate microcentrifuge tubes). Trials were conducted on 12 dates in midsummer (14 June – 1 July 2019).

#### Laboratory Experiment

Field experiments identified DMTS as a chemical that deters use of a carcass for breeding. To examine a possible behavioral mechanism, we assessed whether DMTS leads to rejection of a discovered carcass. Female *N. orbicollis* from a laboratory-reared colony derived from the Bethany population (25–35 days old) were provided a fresh carcass (15–19 g) in a breeding container (35 x 11 x 18 cm) half-filled with commercial topsoil. In half the trials (N = 20), a microcentrifuge tube (punctured with a 26 g needle) was supplied with 10 μl of DMTS on days 1, 3 and 5 so that DMTS would be present at least through the first 8 days of the trial (until all chemical had volatilized). When larvae dispersed from the nest, they were counted and weighed.

#### Statistical Analysis

Field experiments were analyzed in two ways. The frequency of discovery by burying beetles was tested using Fisher’s Exact test (carcasses scavenged by vertebrates excluded). In addition, a score (total number of beetles discovering carcasses for a treatment on a given day, divided by the number of unscavenged carcasses for that treatment) was compared among treatments using a paired test. Each date of sampling, with three traps per treatment, was a single experimental replicate. The mean number of beetles per trap was not normally distributed, contained many zero values and was highly skewed; standard transformations did not result in near-normal distributions. A nonparametric test (Wilcoxon’s Matched Pairs Signed Ranks test) was therefore employed to examine treatment differences in scores (SAS Institute Inc 2007). Trials for experiments 4 and 5 were conducted at two sites (Bethany and Woodbury). Within these experiments, there were no significant differences between sites for either discovery rate or number of beetles per trap; trials from these sites were combined for analysis.

In the laboratory experiment, the probability of breeding in the presence or absence of DMTS was assessed by a Binomial test. The number of larvae, total mass of a brood and mean larval mass were compared using an Analysis of Covariance with treatment the dependent variable and carcass mass the covariate.

## RESULTS

### Experiment 1 – Antimicrobial Treatment of a Carcass

Carcasses aged for two days in the laboratory prior to being placed in the field were more attractive to beetles than fresh carcasses; two-day carcasses treated with an antibacterial were intermediate in attractivity. This was reflected by both discovery rates (all pairwise comparisons p < 0.001; Fisher’s Exact test) and by the mean number of beetles per trap-night (Fig. 2). Antifungal treatment did not affect carcass discovery (Supplementary Material, Fig. S1).

**Fig. 2.**
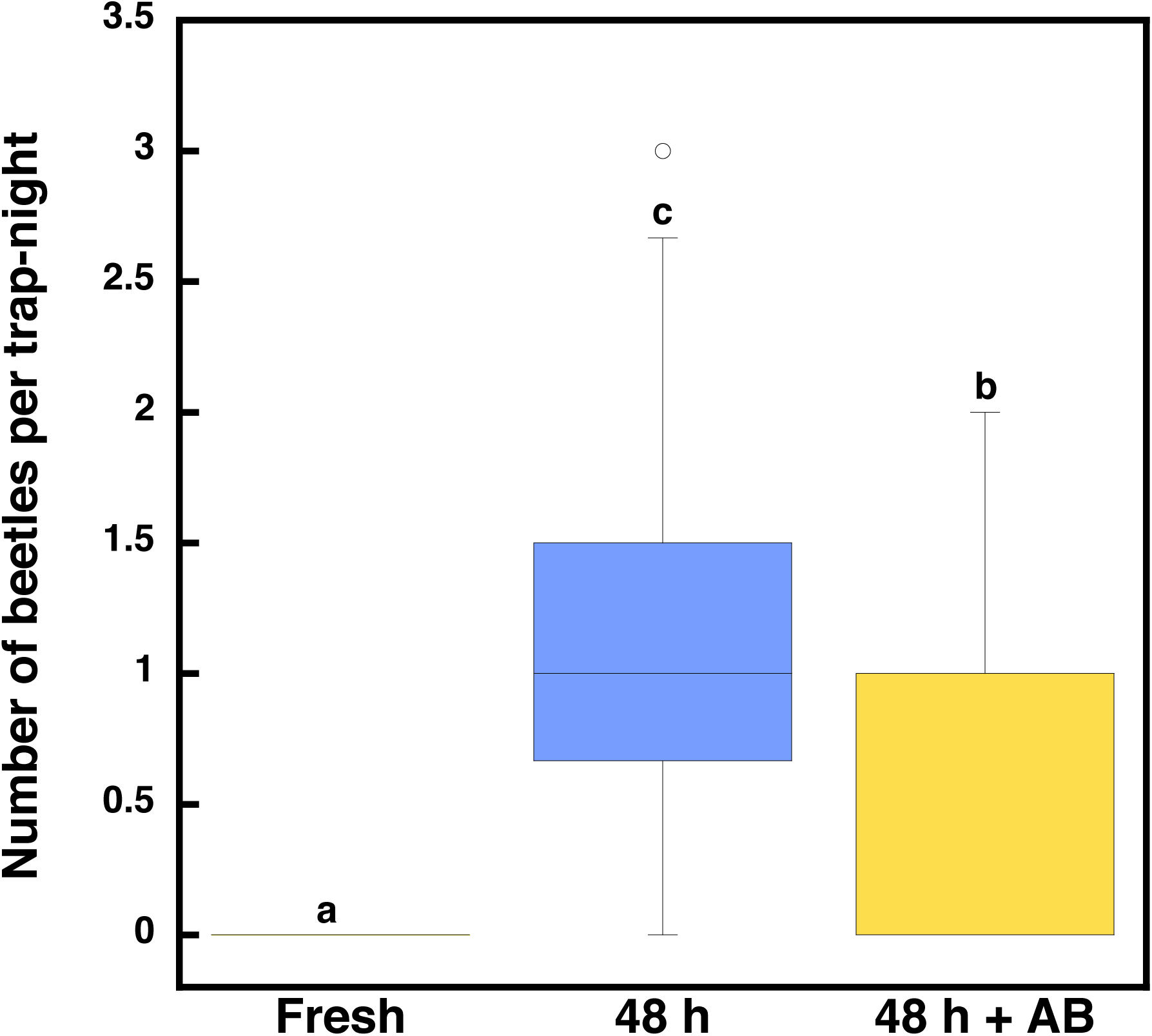
Attraction of burying beetles to fresh carcasses, carcasses aged for 48 h, and carcasses aged for 48 h that were treated with an anti-bacterial (AB). Shown are medians (horizontal lines), the middle quartiles (boxes), and outliers (markers). The upper stem and cap bars represent the upper quartile + 1.5*interquartile distance (SAS Institute Inc 2007). Different letters above the bars indicate significant differences (P < 0.01, Wilcoxon’s Matched Pairs Signed Ranks test)

### Experiments 2-6 – Chemical Supplements

Results for the effects of chemical supplements on the discovery of a fresh carcass are summarized in Table 1. For DMS, the discovery rate and number of beetles attracted per trap were not significantly different for either *N. orbicollis* in midsummer (Exp. 2a) or *N. tomentosus* in late summer (Exp. 2b, Table 1). Combining trials from the two species resulted in significant differences for both measures (discovery rate, P = 0.03, Fisher’s Exact test; number of beetles per trap, P = 0.015, Wilcoxon’s MPSRT). We also found only weak support for DMDS as an attractant (Exp. 3, Table 1; discovery rate, P = 0.017, Fisher’s Exact test; number of beetles per trap, P = 0.11, Wilcoxon’s MPSRT). The presence of DMTS as a supplement reduced the number of beetles finding and burying a fresh carcass — nearly three times as many beetles came to a fresh carcass as one in the presence of DMTS. The difference in discovery rate was also highly significant (Exp. 4, Table 1). The presence of MeSCN as a supplement promoted carcass discovery. The overall rate of discovery for MeSCN carcasses in experiments 5A, 5B and 6 was 93.4%, far higher than any other treatment in any other experiment. The ability of MeSCN to draw beetles to a carcass, however, was diminished in the presence of DMTS, providing additional evidence that DMTS deters breeding *Nicrophorus* from coming to a carcass. The presence of MeSAc did not affect the rate of carcass discovery or the number of beetles found at a fresh carcass (Exps. 5a & 5b, Table 1).

#### Laboratory Experiment

In the laboratory, the presence of the repellent identified from field experiments, DMTS, did not affect the probability that a carcass would produce a brood as all carcasses (N = 20 per treatment) were utilized by *N. orbicollis* (P = 1.00, Fisher’s Exact test). There were also no differences of chemical treatment for number of larvae (DMTS: 14.75 ± 0.81; Control: 15.60 ± 0.67), total brood mass (DMTS: 5.17 ± 0.22 g; Control: 5.49 ± 0.18 g) or mean mass of larvae (DMTS: 0.359 ± 0.010 g; Control: 0.357 ± 0.010 g) and no significant chemical x carcass mass interactions (Table 2).

**Table 2.** Significance test for reproductive output of *N. orbicollis* when breeding in the presence or absence of the chemical volatile DMTS

|  | Number of larvae | Mean mass of larvae | Total brood mass |
| --- | --- | --- | --- |
| Explanatory variable<sup>1</sup> |  |  |  |
| Chemical treatment | $F = 0.58, P = 0.45$ | $F = 0.22, P = 0.89$ | $F = 1.31, P = 0.26$ |
| Carcass mass | $F = 4.74, \mathbf{P = 0.04}$ | $F = 0.02, P = 0.88$ | $F = 14.24, \mathbf{P < 0.001}$ |
| Chem x carcass mass | $F = 0.34, P = 0.56$ | $F = 0.53, P = 0.47$ | $F = 0.34, P = 0.56$ |

<sup>1</sup>All explanatory variables with 1 df

## DISCUSSION

We highlight two findings. MeSCN, which has received little study, has a remarkable ability to attract breeding burying beetles to a fresh carcass and its presence almost guarantees discovery when breeders are highly active. DMTS, known to be an important attractant for both carrion beetles and dipterans (Kalinova et al. 2009; von Hoermann et al. 2016; Yan et al. 2018; Zito et al. 2014), deters *N. orbicollis* from a resource for breeding. These findings, which suggest that insects seeking a fresh carcass are using a different set of cues than most carrion-frequenting insects, are discussed in greater detail below. We note that the two chemicals with the greatest effects were the least volatile, which may evoke responses of insects over greater distances (Brodie et al. 2016; Schlyter et al. 1987).

Antibacterial treatment of a 48 h carcass decreased discovery by burying beetles (Exp. 1), likely because bacterial-derived cues were reduced. We found no evidence that fungi on carrion contribute to the cues for burying beetles (Supplementary Material). The primary cues are sulfur-based, in part, because animal protein is rich in the sulfur-containing amino acids cysteine and methionine that bacteria metabolize. Specific bacteria have been identified that produce DMDS, DMTS, MeSAc and MeSCN (Cammack et al. 2015; Kai et al. 2009; Lam et al. 2010; Ossowicki et al. 2017; Paczkowski et al. 2012).

MeSCN (as thiocyanic acid, methyl ester) has recently been listed as a volatile of carrion, appearing with the first measurement after death (Armstrong et al. 2016). It had not been reported in earlier inventories of carrion volatiles (Dekeirsschieter et al. 2009; Forbes and Carter 2015; Vass et al. 2008). The only behavioral study of MeSCN that we are aware describes its ability to repel the Gram-negative *Pseudomonas aeruginosa* Schröter (Ohga et al. 1993). Along with DMDS, DMTS and MeSAC, MeSCN is produced by the rhizospheric *Psudomonas donghuensis* Gao, possibly as an antifungal agent protecting against potential plant pathogens (Ossowicki et al. 2017). Our study suggests MeSCN may have a previously unrecognized role as an indicator of a recently dead animal. Its exploration may help to fill gaps in our knowledge of the cues used by early colonizers of carrion.

The sulfides, DMS, DMDS and DMTS can also be detected at low levels in the early phase of animal decomposition and are well-recognized cues for numerous carrion insects (Armstrong et al. 2016). Based on differences in the timing of peak production (Kalinova et al. 2009), we had predicted that DMTS would enhance discovery of a carcass for breeding, but less so than DMS and DMDS. DMTS, however, was found to be an outright repellent. Some insects bypass a potential resource because of a marked change in the sensitivity of sensory receptors. The mosquito *Culex pipiens* L., for example, loses the ability to respond to host-specific cues during reproductive diapause (Bowen et al. 1988). This does not appear to be the case here, however, as *N. orbicollis* does not ignore DMTS, but actively avoids it. We speculate that this occurs because high levels of DMTS may indicate an older or flyblown resource that is less optimal for burying beetle reproduction (see Rozen et al. 2008; Trumbo et al. 2016).

Previous studies found that DMDS and especially DMTS were strong attractants for both carrion flies (Brodie et al. 2014; Chaudhury et al. 2015; Frederickx et al. 2012; Nilssen et al. 1996; Zito et al. 2014, but see Lam et al. 2017) and silphid beetles, including *Nicrophorus* spp. (Dekeirsschieter et al. 2013; Kalinova et al. 2009; Podskalska et al. 2009). There are several possible explanations for the different responses reported for breeding burying beetles to DMTS. Beetles on the ground that are walking toward a discovered carcass may respond differently than beetles in flight. *Nicrophorus pustulatus*, for example, has never been found to breed on carrion in the field, but will use this resource if placed directly on it in the laboratory (Smith et al. 2007). This suggests that a burying beetle on the ground will respond to a greater range of resources than while in flight. Kalinova et al. (2009) found that *N. vespilloides* and *N. vespillo* walking in a Y-maze favor DMTS over a blank. This is not surprising as DMTS does not preclude breeding in the laboratory (present study) and once on the ground, it likely is advantageous to assess the suitability of a carcass directly (Trumbo et al. 1995). Interestingly, Rozen et al. (2008) showed in a choice study that *N. vespilloides* Herbst first walks toward a more decomposed mouse carcass rather than a fresh carcass, but then ultimately chooses the fresher resource after inspection.

A burying beetle in flight, however, may be more selective. A volatile cue that indicates that a resource is likely too large, too maggot-infested or too damaged for monopolization by a burying beetle may act as a repellent, allowing the beetle to avoid the energetic cost of searching for an unusable resource. Recinos-Aguilar et al. (2019) found that maggot infestation dramatically accelerates the release of DMTS from a carcass, resulting in higher levels at two days with maggots than at 4 days without maggots (a similar effects of maggots on DMDS release can occur, Chen et al. 2020). Just as feeding burying beetles avoid fresh carcasses (von Hoermann et al. 2013), breeders may avoid flyblown carcasses, at least in flight. If a breeder finds such a carcass, its aggression is reduced (Chen et al. 2020) and it does not exhibit the rapid development of ovaries that occurs with a fresh carcass (Wilson and Knollenberg 1984). A damaged (incised) carcass also releases more DMTS than a same-aged undamaged carcass, even if no maggots are present; dipterans are subsequently attracted to the incised area (Brodie et al. 2014; Brodie et al. 2016). Opened carcasses decay faster (Mann et al. 1990) and are of less value to burying beetles as a resource for breeding, even when maggots are not present (Trumbo 2017).

The field work of Podskalska et al. (2009), where *N. vespillo* L. (but not other burying beetles) came in large numbers to traps baited with DMDS and DMTS, is more difficult to reconcile with the present study. There were differences in the length of the trial and the amount of chemical, but these were minor and are unlikely to explain the difference in response to DMTS. Podskalska et al. (2009) also employed all three chemicals simultaneously, which in combination could have acted as an attractant. We did find that DMS and DMDS could be attractive, but the effect was small and much less than for MeSCN. We found that DMTS did not enhance but rather detracted from the attractiveness of MeSCN. We also provided a carcass for burial to test for breeding behavior rather than use a chemical-only bait. It is unclear why this might switch DMTS from the most attractive chemical in Podskalska et al. (2009) to the most repellent chemical in the present study. Two other factors may be salient. The experiment detailed in Podskalska et al. (2009) occurred in late summer, so it is possible that post-breeding beetles or their newly emerged adult offspring were in reproductive diapause and were feeding in preparation for winter. They would then be seeking carrion in active decay rather than fresh. We have found that *N. pustulatus* Herschel comes in large numbers to traps with DMDS and DMTS (unpublished results). This species will feed on well-rotted carrion and maggots but never has been found to breed on fresh carrion in the field, instead using snake eggs for that purpose (Blouin-Demers and Weatherhead 2000; Smith et al. 2007). A second difference was that the experiment by Podskalska et al. (2009) occurred in an open field, where DMTS may be used differently by carrion insects than in woodlands. More work needs to be done on species and habitat differences, chemical interactions, and post-breeding responses of beetles.

Indirect evidence that DMTS indicates an aged, well-rotted carcass (suitable for feeding but not breeding burying beetles) is supported by work on other uses of this volatile. The earwig, *Labidura riparia* Pallas, uses DMDS and DMTS to defend itself against lizards by mimicking rotting-flesh odor, a trait that would be ineffective if the odor represented fresher carrion (Byers 2015). Stinkhorn fungi, whose common name reflects that it is not mimicking a fresh carcass, use these chemicals to attract flies to disperse their spores (Johnson and Jürgens 2010). At least five independent lineages of corpse-mimicking plants have evolved a bouquet that mimics the strong odor of the active decay stage. DMDS and DMTS have consistently been identified as important in those plants where a volatile analysis has been completed (Borg-Karlson et al. 1994; Jürgens et al. 2013; Kite et al. 1998). It is likely that these plants evolved to mimic the stage that attracts the highest number of deceived pollinators during active decay (see Kočárek 2003) rather than a fresh carcass that would attract a limited number of early colonizers (reviewed in Jürgens and Shuttleworth 2015). MeSAc is also produced by corpse-mimicking plants, but its role has not been explored (Kite and Hetterscheid 2017; Shirasu et al. 2010). We recently uncovered evidence that MeSAc has a powerful enhancer effect as a synergist of DMTS, attracting nonparental silphine beetles that specialize on the active decay stage (Trumbo and Dicapua 2020). Neither MeSAc nor DMTS may be a reliable cue for locating a resource for reproduction by burying beetles and may be ignored or actively avoided during volant searches by breeders.

Ecological succession of insects on a corpse has been known for at least 130 years (Mégnin 1883). The fact that a mostly ignored sulfur chemical is by far the greatest attractant to a carrion insect seeking a fresh carcass, and that a previously identified attractant was the most repellent, suggests that we still have much to learn. Our study suggests that MeSCN should be explored as an attractant for the endangered *N. americanus* Olivier and other rare *Nicrophorus* spp.; a positive finding might greatly facilitate population monitoring and mark-recapture studies. The possible response of forensically-important dipterans to MeSCN is also of interest. Finally, as more information is gathered on the volatiles that attract insects to the various stages of decomposition, it may be possible to construct a mechanistic model of succession on carrion based on the changing profile of microbially-derived cues (Michaud and Moreau 2017; Pechal et al. 2013).

## Author Contributions

ST and SS planned and designed the research. ST carried out the experiments, analyzed the data and wrote the manuscript.

## Acknowledgements

Sebastian Wojcik assisted in the maintenance of beetle colonies. The Southern Connecticut Regional Water Authority, the Flanders Preserve and the State of Connecticut Department of Energy and the Environment granted permission for field experiments. This research was supported by The University of Connecticut Research Foundation (STT). The authors declare no conflicts of interest.

## ONLINE RESOURCE – SUPPLEMENTARY MATERIAL

### METHODS

Five types of carcasses were placed in the field (Bethany site) to investigate the source of cues that free-flying beetles use to locate a resource for breeding. Control carcasses were thawed and aged in the laboratory for 48 h on woodland soil from the field site before placement in the field (see Exp. 1). They were immersed in water at 3h, 24 h and 48 h to standardize handling. Fresh carcasses were thawed at room temperature for 2-6 h and then immersed in water prior to placement in the field. Antibacterial-treated carcasses were immersed in an aqueous tetracycline hydrochloride solution (45 mg per 100ml) (Sigma Aldrich) in a closed container and tumbled to ensure complete topical exposure at 3h, 24 h and 48 h. Antifungal-treated carcasses were immersed in a Nystatin**®** suspension (35 mg per 100ml) and tumbled as above on the same schedule. An antibacterial + antifungal treatment employed both topical applications sequentially. Antibacterial and antifungal solutions were kept refrigerated (4 ^0^C) and used within 48 h to ensure potency.

Fifteen 9-12 g mouse carcasses (3 for each of the 5 treatments) were placed in the field on each of 5 dates in July and 5 dates in August/ September 2014 (total of 150 carcasses). Carcasses were placed on top of soil in a plastic cylinder (10 cm diameter, 12 cm height, filled 4/5 with soil and buried flush with the surface). To minimize cross-attraction between treatments, 3 carcasses, all from the same treatment, were placed on a 3-point transect, with transect points 20 m apart. There were 5 transects, one for each treatment, at a minimum 200 m distance from other transects. Treatments were rotated through the transects such that each transect was used once for each treatment in July and once in August/September. In July, when the most active burying beetle (*N. orbicollis*) was nocturnal, carcasses were placed in the field at 17:00 and checked at 9:00 the following day. In late August/September, when the most active burying beetle (*N. tomentosus*) was diurnal, the carcasses were placed in the field at 10:00 and checked the same day at 18:00. Because the overall discovery rate of carcasses was similar during the two time periods (33.3% and 30.7%, Fisher’s Exact test, P = 0.86), data were combined for analysis. To terminate the trial, the soil contents of each container was examined for the presence and number of burying beetles. All containers were removed from the field after each trial and cleaned before the following trial to remove residual cues. The data were analyzed as for Exp. 1.

### RESULTS

Antibacterial treatment had clear effects on the ability of burying beetles to discover a carcass placed on the ground. Antibacterial treatment reduced discovery of a 2-day carcass from 67.9% (48 h Control) to 15.4% (48 h + AB; 14.3% for 48 h + AB/AF). Antifungal treatment alone had no such effect (77.4%, 48 h + AF). Fresh carcasses (thawed for 2 - 6 h prior to the start of the activity period) were very difficult for burying beetles to locate (3.8%). The mean number of beetles per trap-night followed a similar pattern to discovery rate (Fig. S1).

**Fig. S1.**
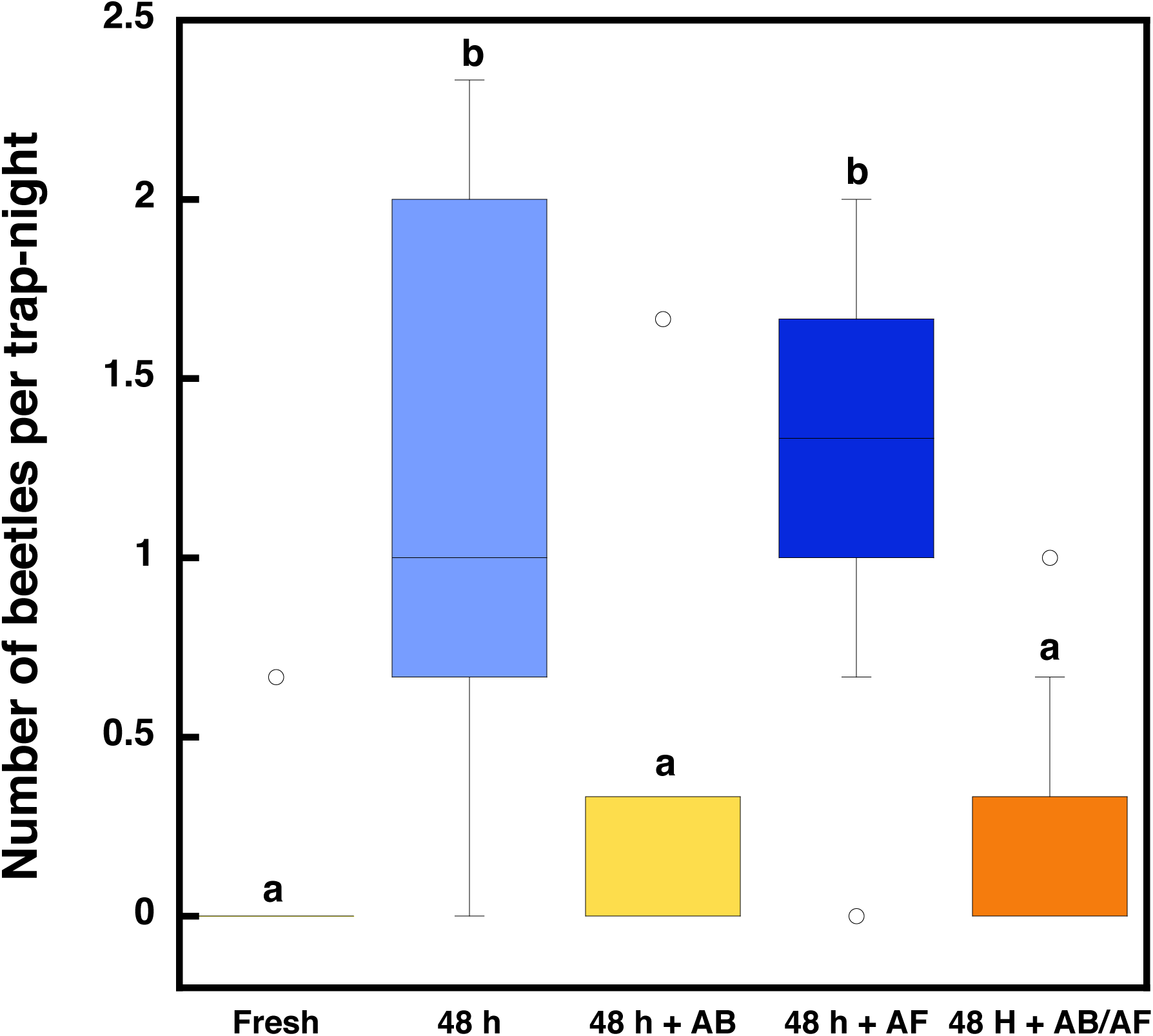
The number of burying beetles per trap-night caught at carcasses that were fresh, 48 h, 48 h + anti-bacterial (AB), 48 h + antifungal (AF) and 48 h + AB/AF. Shown are medians (horizontal lines), the middle quartiles (boxes), and outliers (markers). The upper stem and cap bars represent the upper quartile + 1.5*interquartile distance. Different letters above the bars indicate significant differences (P < 0.01, Wilcoxon’s Matched Pairs Signed Ranks test)

